# SigTools: Exploratory Visualization for Genomic Signals

**DOI:** 10.1101/2021.08.02.454408

**Authors:** Shohre Masoumi, Maxwell W. Libbrecht, Kay C. Wiese

**Affiliations:** Simon Fraser University, School of Computing Science

## Abstract

**Motivation:** With the advancement of sequencing technologies, genomic data sets are constantly being expanded by high volumes of different data types. One recently introduced data type in genomic science is genomic signals, which are usually short-read coverage measurements over the genome. An example of genomic signals is Epigenomic marks which are utilized to locate functional and nonfunctional elements in genome annotation studies. To understand and evaluate the results of such studies, one needs to understand and analyze the characteristics of the input data.

**Results:** SigTools is an R-based genomic signals visualization package developed with two objectives: 1) to facilitate genomic signals exploration in order to uncover insights for later model training, refinement, and development by including *distribution* and *autocorrelation* plots. 2) to enable genomic signals interpretation by including *correlation*, and *aggregation* plots. Moreover, Sigtools also provides text-based descriptive statistics of the given signals which can be practical when developing and evaluating learning models. We also include results from 2 case studies. The first examines several previously studied genomic signals called histone modifications. This use case demonstrates how SigTools can be beneficial for satisfying scientists’ curiosity in exploring and establishing recognized datasets. The second use case examines a dataset of novel chromatin state features which are novel genomic signals generated by a learning model. This use case demonstrates how SigTools can assist in exploring the characteristics and behavior of novel signals towards their interpretation. In addition, our corresponding web application, SigTools-Shiny, extends the accessibility scope of these modules to people who are more comfortable working with graphical user interfaces instead of command-line tools.

**Availability:** SigTools source code, installation guide, and manual is available on http://github.com/shohre73.

## 1 Introduction

Understanding the structure, behavior, and interactions of the DNA contents of organisms is the fundamental motivation of Genomics science. The hypotheses in genomics heavily rely on voluminous and diverse data extracted from the organism’s cell. Moreover, with the advancement of sequencing technologies, genomic data is being generated at a high pace, making the analysis phase the bottleneck of genomic studies (Navarro *et al.*, 2019).

A genomic signal is a continuous variable across the genome indicating the presence of a biological activity such as protein interaction, DNA methylation, transcription sites, chromatin crosslinks, and regulatory elements (ENCODE Project Consortium, 2012). To obtain a signal in a lab, a high-throughput sequencing device *selectively* sequences DNA fragments which are then mapped back to the original genome.

Genomic coordinates associated with the coverage measurements of this mapping is recorded as a genomic signal. RNA-Seq, ChIP-seq and ATAC-seq are technologies specifically designed to generate transcriptome, interactional, and chromatin accessibility signals respectively. These signals are utilized in detecting the position of different elements on the genome and discovering the reason why different cell types emerge from the same DNA material (Hoffman *et al.*, 2012a; Jason and Manolis, 2017), investigating gene regulation in cancer research (Gul, 2017), and understanding 3D genome architecture (Mishra and Hawkins, 2017). Hence, several visualization tools have been developed to leverage the human visual system in explorations of genomic signals behaviors and characteristics.

Many of these tools, namely *genome browsers* (e.g. WJ *et al.*, 2002; Buels *et al.*, 2016; Zhou *et al.*, 2011), preserve the sequential nature of these signals by presenting them in a linear or circular layout, possibly with parallel arrangements to enable comparison between signals (Nusrat *et al.*, 2019). Such tools are commonly used to investigate local behaviors around specific regions, for example, to depict regulatory elements near a particular gene (Gosselin *et al.*, 2017) or displaying read numbers for different signals at a specific locus (Sabari *et al.*, 2018).

Other tools employ statistical procedures to illustrate the global behavior of the signals. Accordingly, this class of tools usually work with multi-sequence alignment data formats (SAM/BAM), rather than continuous-valued data formats (WIG/bigWig/BED/bedGraph) which contain the actual value of the signals at each position or bin.

deepTools (Ramírez *et al.*, 2016) offers several modules for pattern discovery and comparison of signal coverage and enrichment over multiple genomic regions. Depicting DNA methylation average values over protein-coding genes to investigate their role in organism development is an example application of deepTools (Daccord *et al.*, 2017). In addition to average coverage plots, ngs.plot (Shen *et al.*, 2014) also includes a database of genomic elements to facilitate region selection.

Segtools (Buske *et al.*, 2011) promotes genomic signals analysis over probabilistic generated genomic elements. As an example, it can be employed to generate a heatmap of histone modification and obtained genomic labels (Hoffman *et al.*, 2012b). Furthermore, it encourages short-read independent signal analysis by using genomedata (Hoffman *et al.*, 2010) as the input format for genomic signals.

Although the reviewed tools above offer a wide selection of processing and visualization features for genomic signals analysis, we identified the need for a set of tools that assists scientists in the early steps of genomic signal analysis, such as the value range, variation, and covariation. Novel genomic signals are being generated both in wet labs by biologists or via learning models by computer scientists. When such researchers come across a new signal, some common questions need to be discussed before this signal could be introduced to the genomic society and be employed in other studies: What numeric range does this signal cover? Does it contain noise? If positive, how much noise is there in the data? How much data variation does it have? Does it behave similarly to any previously studied signal? How does it behave in general around specific genomic elements?

The objective of this project is to provide a cohesive package that contains all the essential data analysis tasks for scientists who are working with novel genomic signals and do not want to carry out additional coding for their project. Furthermore, we also introduce SigTools-Shiny, a web-based graphical user interface that includes all SigTools preprocessing and visual modules as an alternative for users who want to eliminate command-line interaction in their experience.

The following section, Section 2, reviews SigTools preprocessing modules. The next section, Section 3, describes SigTools visualization modules. Section 4 is dedicated to the Shiny App interface, data flow, and interactivity. Section 5 includes two use cases to demonstrate SigTools utility. The first use case examines several previously studied genomic signals named histone modifications. This use case demonstrates how SigTools can be beneficial for satisfying scientists’ curiosity in exploring and establishing recognized datasets. The second use case examines a dataset of novel chromatin state features which are novel genomic signals generated by a learning model (Chen *et al.*, 2019). This use case demonstrates how SigTools can assist in exploring the characteristics and behavior of novel signals towards their interpretation.

## 2 Data and Preprocessing Modules

Sequenced reads constitute the underlying data for genome-related studies. To *align* these reads is to identify their overlaps, and it is the principal method for reducing the size and complexity of their datasets. Mapping these reads back to a reference sequence results in distinct coverage measurements across its bases. Rather than aiming for uniform coverage, some sequencing methods such as RNA-Seq, ChIP-seq, and ATAC-seq, target specific DNA fragments that are bound to certain proteins and generate a mass of short reads from these isolated regions. A *genomic signal* contains the exact or normalized values of such coverage (Anders and Huber, 2010; Bayat and Libbrecht, 2020).

Apart from visualization modules, SigTools includes the following modules to process and perform optional changes to the available data:

- `sigtools_convertToMultiColBedg`: this module converts several bedGraph files with different bin sizes to a single `mulColBedg` file at the desired resolution. Changing the resolution of genomic signals data sets to a larger bin size is a beneficial strategy for reducing data points particularly for visualizations that discuss an entire chromosome or genome.
- `sigtools_sampling`: besides increasing signals resolution, choosing random stretches of signals is a technique that can be employed to reduce signals’ file size. This approach is particularly useful for obtaining quick results.
- `sigtools_concatenation`: for a particular cell type, the signal data for different chromosomes is usually stored in separate files. This module appends multiple input files together and outputs a single large `.mulColBedg` file that can be used for whole-genome or multiple cell-type analysis. `sigtools_sampling` can optionally be incorporated into this process, preventing the final file to become too large.
- `sigtools_stats`: this module outputs a `.csv` containing the five-number summary (min, lower quartile, median, mean, upper quartile, max) of the present signals in a `.mulColBedg` file. Although SigTools focuses on visual analytics, the text-based output of this module is beneficial in pipeline and learning model development.

## 3 Visualization Modules

Exploratory Data Analysis (EDA) is an essential step in any data-dependent study that highlights data anomalies, patterns, and provides a deeper understanding of the data. SigTools visualization modules (listed below) can be employed to explore the range, shape, variation, and covariation of this data.

Genomic signals data sets generally contain a large population of zeros. Hence, the mean value of a signal is a small number close to zero. *Blacklist* (Amemiya *et al.*, 2019) genomic regions have empirically shown to only commit artifact data in next-generation sequencing. Accordingly, the dismissal of these regions has proved to improve the result of several genomic signal related studies. In SigTools, whenever the `enrichment` parameter is set to be `TRUE` for a signal, given a blacklist input file, the blacklist regions would be excluded, hence eliminating the bias that the large zero population or redundant extreme values introduce to the mean value, and then the input signal would be normalized by its mean value.

### 3.1 sigtools_distribution

This module offers three recognized distribution plots for depicting value frequency: empirical cumulative distribution, kernel density plot, and boxplot. Depicting the value frequency of a variable is a quick approach to get an estimation of its primary characteristics; namely the existence of multiple local maxima, the overall range, and the outliers. Prominent parameters for this module are:

- `percentile`: users can specify what percentile of the data they want to work with. [default:`100`]
- `nozero`: if set to `TRUE`, zero entries would be removed from the data.
- `enrichment`: if set to `TRUE`, the plots would be generated with the enriched values.

### 3.2 sigtools_autocorrelation

The autocorrelation plot is an orthogonal plot with x-axis presenting lags (shift), and y-axis presenting the value of autocorrelation at each lag, which is the correlation of the signal with itself when shifted *lag* times.

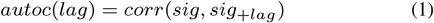

`sigtools_autocorrelation` module generates a line plot of the input signals’ autocorrelation. For each signal, this plot provides a measure of the dependency among consecutive bins or adjacent elements such as neighboring genes (Deng *et al.*, 2010). Signals with higher autocorrelation have smoother picks and valleys. A signal with little variation indicates smaller active regions. The prominent parameters here is `mode`. If set to “`regions`”, another interval file would be required and the autocorrelation would be calculated over adjacent specified regions.

### 3.3 sigtools_correlation

This module outputs a heat-map presenting the pair-wise correspondence of two sets of different signals. A high correlation coefficient between two signals indicates a high behavioral similarity in the sense that wherever the value of one of the signals increases, an increase in the other signal’s value should be observed.

### 3.4 sigtools_aggregation

This module illustrates the overall behavior of a signal over recurring genomic elements. Answering questions such as “Does the signal have a high value within gene regions?” “How does the signal behave over enhancer regions?” “Does the signal have a distinguished behavior over genes upstream or downstream?” could be most helpful for the signal’s interpretation relative positions.

The primary information of the elements under examination —such as the chromosome they are located on, their starting and ending coordinate, and direction— should be supplied as a `.bed` or `.gene_info` file. For a signal *S*, `sigtools_aggregation` computes an aggregation matrix. For a specified element, an array of *S*’s value over that element is retrieved. As the input elements vary in length, these arrays do not have the same length, so unless they undergo an operation that unifies their length, they can not be assembled into a matrix. The `mode` parameter is the prominent parameter for this module which controls the length unification process. If set to `point`, all retrieved arrays would be centrally aligned. However, if set to `region`, the arrays’ length would be unified by smooth interpolation.

## 4 SigTools-Shiny Web Application

Command-line packages are necessary for pipeline development yet a large number of prompt parameters often discourage users to explore their data. Graphical user interfaces are ideal choices for users with limited command-line experiences to interact with their data. Shiny is an R package enabling the assembly of an interactive web-based user interface from R scripts (RStudio, Inc, 2013). The increasing number of Shiny apps in data visualization, particularly genomic data visualization (Yu *et al.*, 2017; Khan and Mathelier, 2017; Reyes *et al.*, 2019; Yu *et al.*, 2019), indicates the effectiveness of this approach in enhancing the accessibility of developed packages. All SigTools utilities are also embodied in an interactive Shiny web application. Along with font size, image size, and axis labels, and other figure customization options, easy change of data parameters in Shiny App facilitates data exploration, selection, and filtering.

SigTools-Shiny is structured based on a Shiny *dashboard* which consists of a vertical navigation bar and a main body. The navigation bar provides access to SigTools-Shiny’s four pages: *Data*, *Plots-static*, *Plots-Interactive*, and *About*. By clicking on each of these sidebar items, their corresponding page appears in the main body.

The *Data* page contains several boxes, one for each data operation discussed in Section 2 and one for data import controls. When a file is uploaded, the five-number summary of its contents appears in a box on the right side of the window, confirming that the uploading process was successful. This summary file can be downloaded using the `Download` button.

The *Plots-Static* page contains only one box named *Canvas* which contains several tabs each for a specific plot: *Boxplot*, *Impirical Cummulative Distribution*, *Kernel Density Distribution*, *Autocorrelation*, *Correlation*, and *Aggregation*. Each tab has a sidebar and a main panel. All the data and plot modification options are located on the sidebar panel. The `GO!` button initiates the plot generation process and eventually the plots is displayed on the main panel.

Much like the *Plot - Static* page, the *Plot - Interactive* page also consists of only one canvas, with multiple tabs. Yet, the plots in this page offer interactions such as zooming in and out and data selection. These plots are generated by Plotly, an R package that enables creating interactive web-based figures.

## 5 Results

The two case studies presented in this section demonstrate SigTools© effectiveness in the interpretation and evaluation of genomic signals data sets. The first case study explores several previously studied genomic signals and concludes that SigTools accurately captures their characteristics. The second case study investigates the attributes of a novel set of genomic signals and exhibits how SigTools can reveal their associate biological function.

### 5.1 Case Study 1 – exploratory analysis of known histone modifications

Histones are proteins that play a crucial role in DNA structure and chromosome organization. It is possible for these proteins to be modified by connecting to a methyl group (methylation) or acetyl group (acetylation) and these modifications heavily impact gene expression. Accordingly, many studies have been focused on histone modification and what particular activity they represent (Bannister and Kouzarides, 2011).

Our first dataset contains six modification signals of protein histone H3 –*H3K4me1*, *H3K4me3*, *H3K9me3*, *H3K27ac*, *H3K27me3*, and *H3K36me3*– over chromosome 21 of the human genome. Table 1 displays the elements and their associated locations that each of these signals represents (Kundaje *et al.*, 2015; Creyghton *et al.*, 2010; Rada-Iglesias *et al.*, 2011). The signals were downloaded from Roadmap Epigenomic Data Portal (Kundaje *et al.*, 2015) in indexed binary format (bigWig) with single base-pair resolution. These files were processed into a multi-column bedGraph file with 200bp resolution using `sigtools_convertToMultiColBedg` function.

**Table 1.** The function and the location of the signals investigated in the first case study.

| Assay | Location |
| --- | --- |
| H3K4me1 | Spatially close to the promoter, though it might lineary be away from the gene. |
| H3K4me3 | Found near the beginning of the gene. |
| H3K9me3 | Contains very few genes. |
| H3K27ac | Commonly found near the transcription start site (TSS). |
| H3K27me3 | Found near the beginning of the gene. |
| H3K36me3 | Gene bodies |

The following mentioned analyses are subfigures of Fig 2, and in this case, they were proposed to explore and acknowledge the already established insights.

**Fig. 1:**
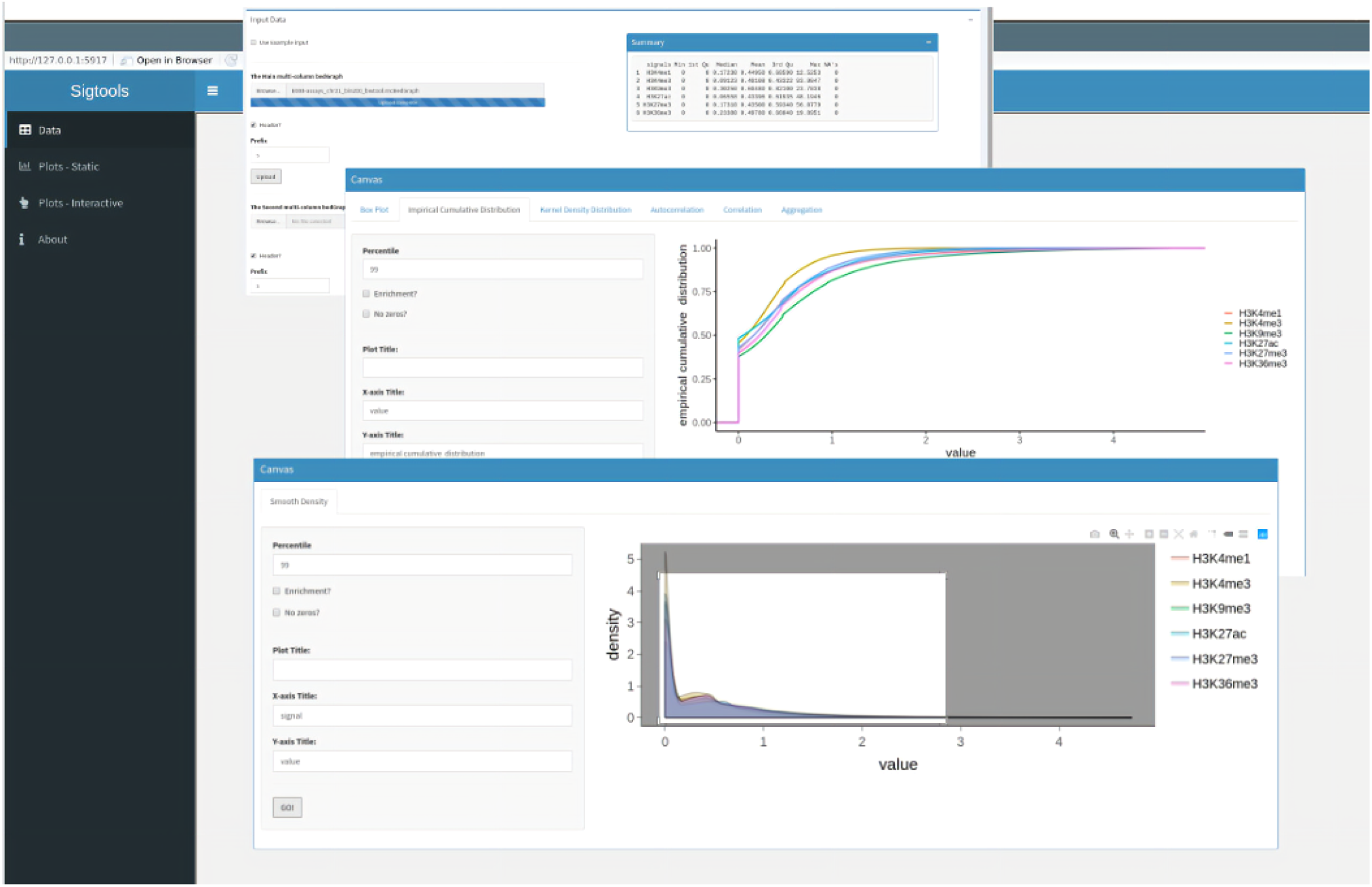
Screen-shots of the main tabs of SigTools Shiny App

**Fig. 2:**
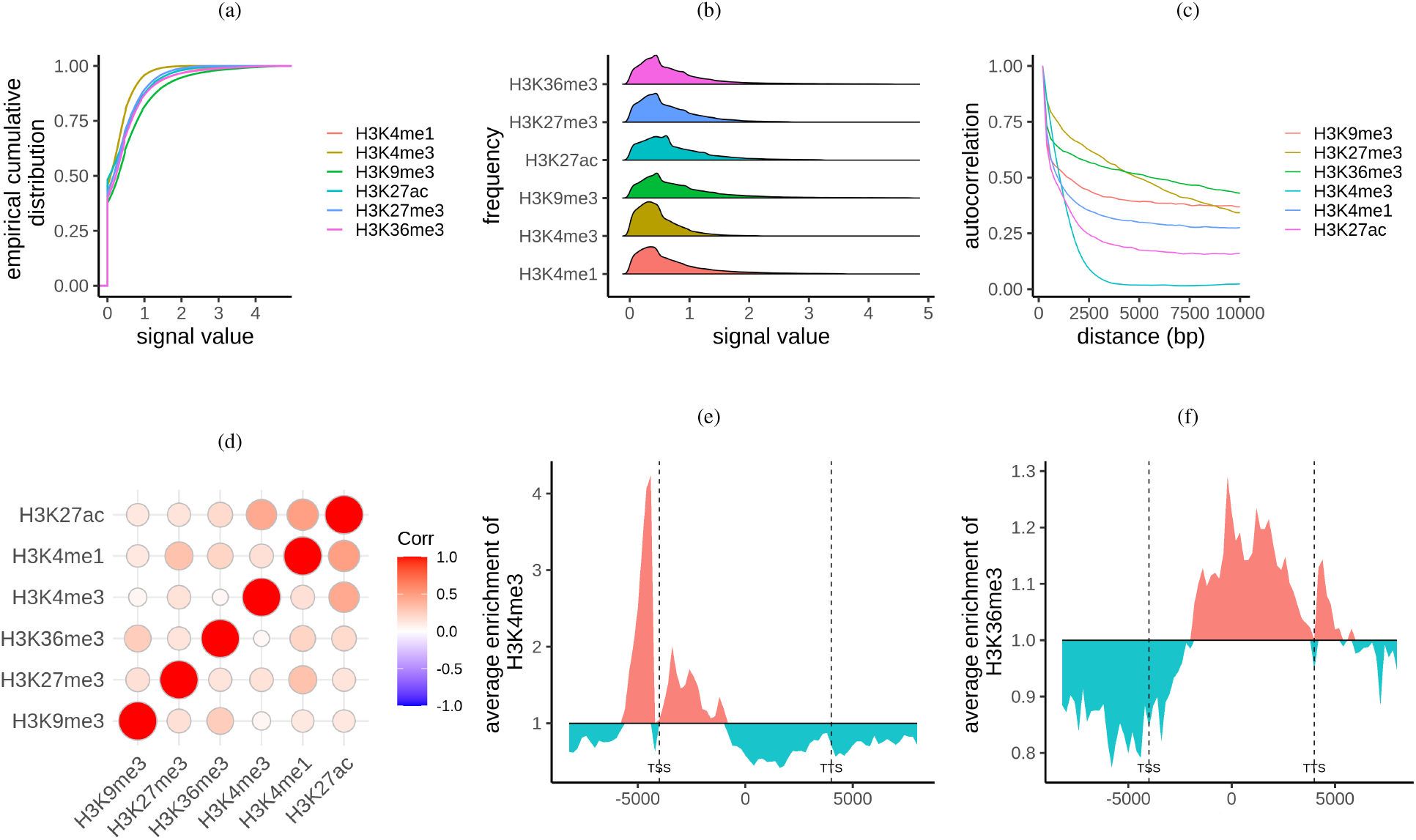
Exploratory analysis of histone modifications: H3K4me1, H3K4me3, H3K9me3, H3K27ac, H3K27me3, and H3K36me3 a) Empirical Cumulative Distribution generated by `sigtools_distribution` with the `plot` parameter set to `ecdf` and `percentile` set to `99`. A high population of zeros is detected. b) Kernel Density Distribution generated by `sigtools_distribution` with the `plot` parameter set to `curve`, `percentile` set to `99`, and `nozeros` set to `TRUE`. It demonstrates the shape and statistical range of the data. c) Autocorrelation plot generated by `sigtools_autocorrealtion` with the `lag` parameter set to `50`. The input file resolution is `200`. This plot indicated that signal H3K4me3 has the most drastic changes among the six discussed signals here. d) Correlation heat-map generated by `sigtools_correaltion` e) H3K4me3 aggregation over chromosome 21 gene regions. Generated by `sigtools_aggregation` f) H3K36me3 aggregation over chromosome 21 gene regions. Generated by `sigtools_aggregation`.

To obtain a quick grasp of the data distribution, we generated an ECDF plot using the 99 percentile of the data (Fig 2a) using `sigtools_distribution` function with `ecdf` option. We observe that repeated zero values constitute almost 50 percent of all the signals, which is not surprising since a great portion of human DNA is non-coding sequences with largely unknown functionality, though some contain regulatory elements (Anandakumar *et al.*, 2017). Next, we examine non-zero values within the 99 percentile by generating a kernel density plot (Fig 2b, `sigtools_distribution` function with `curve` option) which uncovers that most of the remaining population rest within the (0, 2) interval. Having an estimation of variable ranges is particularly necessary when deciding whether to apply any normalization techniques on data before directing it to a learning algorithm.

Having studied value variation, the autocorrelation plot indicates how sudden or smooth the values shift in consecutive bins. Fig 2c displays that out of the six modifications, H3K4me3 has the sharpest picks and deepest valleys, hence it has the smallest active regions. Accordingly, the signal with the highest autocorrelation, H3K36me3, has the largest active regions since not much value transformation is indicated.

To understand if there is a linear association between these value variations, we generated the correlation plot, which uses two visual variables (size and color) to encode Pearson correlation for all pairs of given signals. In this case, Figure 2d displays a high correlation between H3K27ac –indicator of active enhancers– and H3K4me1 –a signal representing all enhancers. Moreover, H3K27ac also demonstrates a high correlation with H3k4me3 which is in accordance with the possible overlap of enhancers and promoters.

The remaining subplots of Fig 2 discuss the average enriched behavior of two of the mentioned histone modifications over gene bodies of chromosome 21. H3K4me3 exhibits high values near the beginning of genes as a promoter does (Fig 2e), and H3K36me3 shows high activity over gene bodies (Fig 2f).

### 5.2 Case Study 2 – interpreting chromatin state feature

A recent study (Chen *et al.*, 2019) proposes chromatin state features for capturing genomic elements instead of discrete annotation. These features are continuous genomic signals which are obtained from histone modifications refined by a Kalman filter state-space model. We chose a set of three features for this case study to demonstrate how SigTools can assist in interpretation of novel genomic signals.

Since these features are to project characteristics of histone modifications, it is expected when the ECDF plot (Fig 3a) of the 99 percentile of feature data displays a large population of zeros in the dataset. We can also obtain an estimation about the range of the data which is about [0, 0.5) for all the signals.

**Fig. 3:**
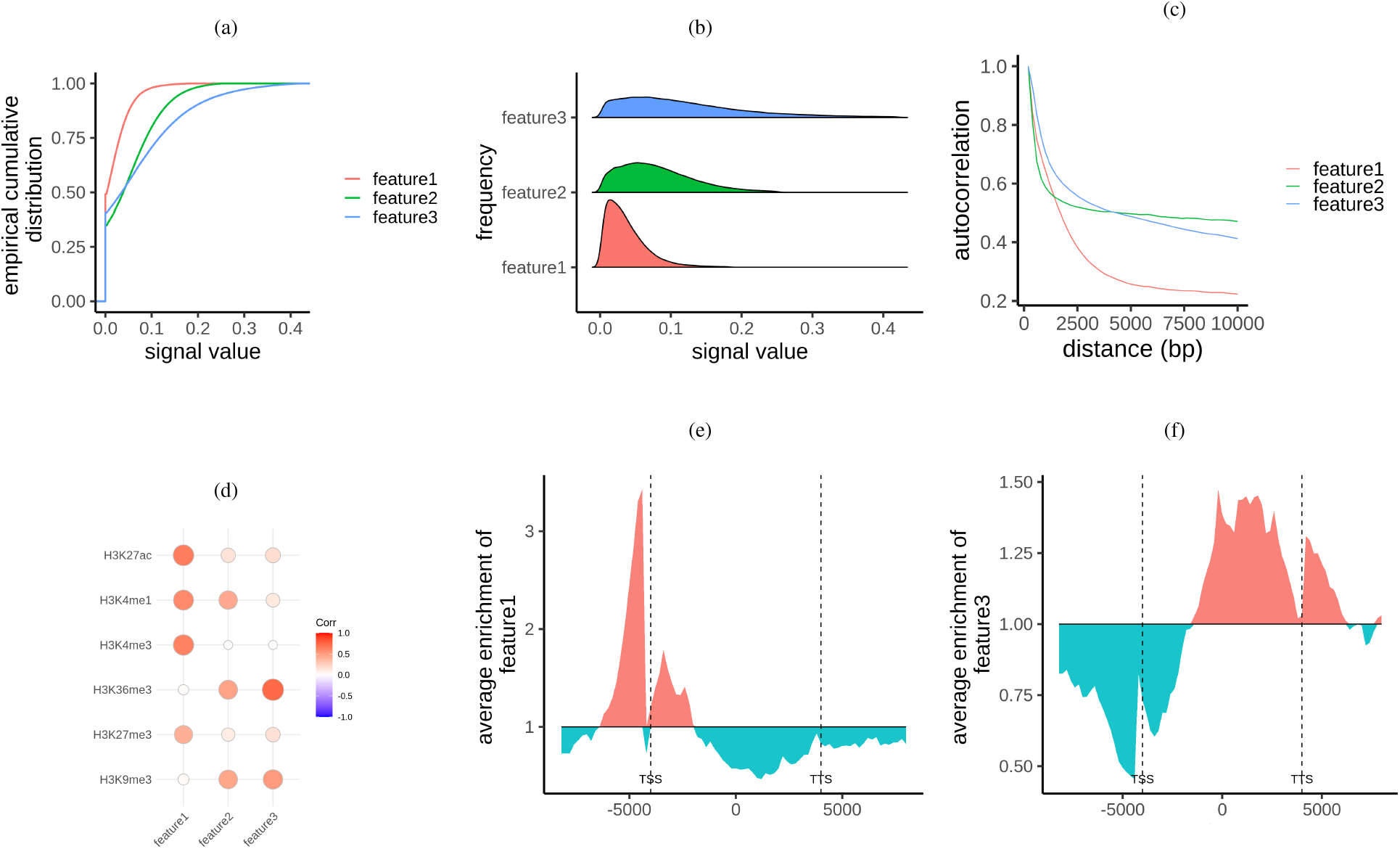
Towards the interpretation of Chromatin State a) Empirical Cumulative Distribution generated by `sigtools_distribution` with the `plot` parameter set to `ecdf` and `percentile` set to `99`. b) Kernel Density Distribution generated by `sigtools_distribution` with the `plot` parameter set to `curve`, `percentile` set to `99`, and `nozeros` set to `TRUE`. c) Autocorrelation plot generated by `sigtools_autocorrealtion` with the `lag` parameter set to `50`. The input file resolution is `200`. d) Correlation heat-map generated by `sigtools_correaltion` e) Aggregation plot of feature1 over chromosome 21 gene bodies. Generated by `sigtools_aggregation` f) Aggregation plot of feature3 over chromosome 21 gene bodies. Generated by `sigtools_aggregation`.

Removing the zero population and the distribution curve plot (Fig 3b) displays that out of the three features, feature1 is denser within the range of (0, 1). Despite this smaller variation, the autocorrelation plot (Fig 3c) displays that feature1 contains more sudden changes.

Fig 3d displays that feature1 mainly correlates with H3K27ac, H3K4me1 and H3K4me3 which are responsible for transcription enhancement. Accordingly, these three assays drop the most in autocorrelation. Plotting the average enrichment of this feature over gene body regions (Fig 3e) make an even stronger argument that feature1 also represents enhancer activity.

## 6 Discussion and Conclusion

An ever-increasing number of genomic signals are being generated by next-generation sequencing technologies and are being widely utilized in studies such as genome annotation (ENCODE Project Consortium, 2012; Hoffman *et al.*, 2012a), cell development, and gene regulation. The analysis of these signals is often associated with the analysis of short-read sequences or genomic elements. However, in recent annotation studies (Chen *et al.*, 2019) theoretically generated signals have been promoted to be a direct representative of biological activities. Regarding the importance and the increasing number of genomic signals and their novel applications we believe there is a growing need for refined tools that enable convenient exploratory analysis and facilitate genomic signals’ interpretation and assessment.

The primary contributions of this work are listed below:

- The design, development, and implementation of an R-based data analysis package, SigTools, to be used by biologists or computer scientists who work with known and novel genomic signals. This package includes several recognized statistical plots that are frequently employed for exploratory data analysis in genomics and other fields. Table 2 is an overview of SigTools visualization modules and their availability in other genomic signal analysis tools. This table indicates that no other tool offers a function for generating distribution and correlation plots for this type of data, the correlation plot is offered only by one other tool, and the aggregation plot is offered by all of them.

**Table 2.**
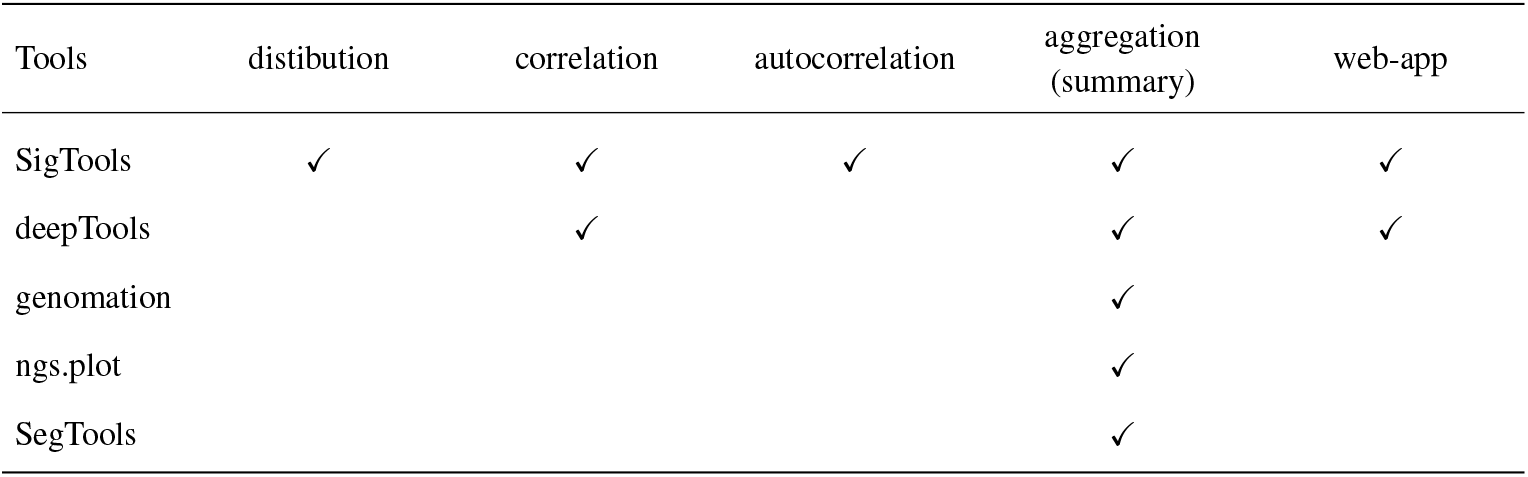
SigTools’ modules and their availability in other tools.

- An aggregation plot is a powerful visualization that has been frequently employed in genomic signals analysis (Fidel Ramírez, 2016; Ramírez *et al.*, 2016; Akalin *et al.*, 2014). Table 2 displays that the aggregation plot is incorporated in all locus-agnostic visualization tools discussed in Section 1. In this work, we *implement* a novel visual encoding for this plot. By introducing a *shifted origin* line to this plot, we aimed to highlight the difference between high and low signal values and enable comparison between different aggregation plots for one signal over different sets of elements, or multiple signals across the same elements.
- Offering a web-based application, SigTools-Shiny. This graphical user interface would be an additional option for users who prefer to limit their interaction with a command-line environment and feel more comfortable inspecting their data with different combinations of plots and parameters through a GUI. SigTools-Shiny also includes some interactive versions of SigTools visualizations, these interactive Java-Script plots are generated by an R package named *Plotly*.

SigTools enables users who work with both experimental or statistical generated genomic signals to obtain text-based or graphical statistical summary of their datasets, to understand what activities their novel signals represent, and investigate the relation of their recently obtained signals with previously studied signals.

As for working with any other extensive dataset, the large size of genomic signals is a challenge that requires close consideration. For obtaining faster results, SigTools offers two solutions: working with modified data to a bigger resolution size, or working with a random subset of the data. The file size can particularly cause issues in web applications when users have to pause their analysis due to multiple uploads when working with diverse datasets. To overcome this issue, some frameworks such as GALAXY (Afgan *et al.*, 2018) offer cloud workstations to their users, hence uploaded data is stored in the user’s account and it can be accessed at any time. As a part of GALAXY, deepTools users benefit from such an online work station. Being a stand-alone tool allowed SigTools to have a flexible user interface design, yet finding a solution for reducing the number of uploads should be included in SigTools future versions. Future versions of SigTools will focus on including an additional number of visualizations and enable comparison between continuous and discrete genomic data.

## Notes

### Competing Interest Statement

The authors have declared no competing interest.

